# Lysosomal Exocytosis Releases Pathogenic α-Synuclein Species from Neurons

**DOI:** 10.1101/2021.04.10.439302

**Authors:** Ying Xue Xie, Nima N. Naseri, Jasmine Fels, Parinati Kharel, Yoonmi Na, Jacqueline Burré, Manu Sharma

## Abstract

Considerable evidence supports the release of pathogenic aggregates of the neuronal protein α-Synuclein (αSyn) into the extracellular space. While this release is proposed to instigate the neuron-to-neuron transmission and spread of αSyn pathology in synucleinopathies including Parkinson’s disease, the molecular-cellular mechanism(s) remain unclear. Here we show that pathogenic species of αSyn accumulate within neuronal lysosomes in mouse brains and primary neurons. We then find that neurons release these pathogenic αSyn species via SNARE-dependent lysosomal exocytosis; proposing a central mechanism for exocytosis of aggregated and degradation-resistant proteins from neurons.

## INTRODUCTION

Cytoplasmic aggregates of the synaptic protein αSyn are a characteristic feature of multiple neurodegenerative diseases termed “synucleinopathies”, including Parkinson disease (PD). Over the course of these age-linked diseases, αSyn assembles into amyloid-type β-sheet rich aggregates, depositing as Lewy bodies and/or Lewy neurites within neurons (Baba et al., 1998; Spillantini et al., 1998; Spillantini et al., 1997). The neuro-anatomical propagation of αSyn pathology is highly stereotyped, and has been defined well-enough to be used in staging PD – originating in the medulla and olfactory bulb, then advancing to the midbrain and basal forebrain, followed by “spread” to the neocortex (Braak et al., 2002; Braak et al., 2003). This observation, combined with detection of extracellular αSyn in human cerebrospinal fluid (El-Agnaf et al., 2003), led to the original conjecture that pathogenic species of αSyn may be transmitted neuron-to-neuron via prion-like permissive templating. Subsequently, in PD patients who had received fetal neuron transplants, appearance of Lewy pathology in the undiseased grafted neurons also pointed to the transmission of αSyn pathology from the surrounding PD-affected neurons (Kordower et al., 2008; Li et al., 2008). These observations were followed by animal studies which confirmed either neuron-to-neuron transmission of αSyn pathology (Desplats et al., 2009), or induction of intraneuronal αSyn pathology by an inoculum of extracellular αSyn fibrils (Luk et al., 2012; Volpicelli-Daley et al., 2011a).

However, unclarity persists in how the cytosolic aggregates are conveyed into the extracellular milieu: Proposed pathway(s) include secretion of αSyn monomers, oligomers and/or larger aggregates inside extracellular vesicles – such as exosomes (Danzer et al., 2012; Emmanouilidou et al., 2010) or microvesicles (Minakaki et al., 2018) – which have been categorized collectively as “unconventional secretion”. Other proposed mechanisms include synaptic vesicle release (Yamada and Iwatsubo, 2018), or exocytosis of other types of vesicles (Jang et al., 2010; Lee et al., 2005), exophagy (Ejlerskov et al., 2013), membrane translocation (Ahn et al., 2006), or trafficking of aggregates through tunneling nanotubes (Abounit et al., 2016). Yet, all of these studies have not identified a neuronal organelle that accumulates and then secretes the pathogenic αSyn aggregates, resulting in the current mechanistic gap.

Here, we present our finding that pathogenic αSyn species accumulate within the neuronal lysosomes in mouse brains and in primary neurons, and that these species are released from neurons via SNARE-dependent lysosomal exocytosis.

## RESULTS

### Pathologenic species of αSyn accumulate within the lysosomes of transgenic αSyn^A53T^ mouse brains

Rapid neurodegeneration and synapse loss in cysteine string protein-α knockout mice (CSPα^−/−^) is rescued by transgenic expression of αSyn carrying the PD mutation A53T (Tg-αSyn^A53T^) (Chandra et al., 2005). However, beyond 5 months, the rescued Tg-αSyn^A53T^/CSPα^−/−^ mice begin exhibiting neurodegeneration due to the transgenic overexpression of αSyn^A53T^. Curiously, we found that Tg-αSyn^A53T^/CSPα^−/−^ mice have accelerated accumulation of the pathogenic versions of αSyn compared to Tg-αSyn^A53T^/CSPα^+/−^ mice, detectable by phospho-Ser129 specific, filamentous αSyn-specific, and amyloid-specific antibodies (**Supplementary Fig. S1**). CSPα^+/−^ mice have normal CSPα function (Fernández-Chacón et al., 2004), and there was no change in monomeric αSyn levels (**Supplementary Fig. S1**). This suggested that brains of Tg-αSyn^A53T^/CSPα^−/−^ mice are collecting αSyn aggregates faster in the absence of CSPα.

Loss-of-function mutations in CSPα cause the lysosomal storage disorder adult-onset neuronal ceroid lipofuscinosis (ANCL) or Kufs disease (Benitez et al., 2011; Cadieux-Dion et al., 2013; Nosková et al., 2011; Velinov et al., 2012). Specifically, loss of CSPα function leads to progressive accumulation of lysosomes containing undegraded material such as lipofuscin/residual bodies, in ANCL patient brains (Benitez et al., 2011), in CSP^−/−^ mouse brains (Fernández-Chacón et al., 2004), in Drosophila models carrying the ANCL mutations (Imler et al., 2019), and in ANCL patient-derived induced neurons (Naseri et al., 2020). Tg-αSyn^A53T^/CSPα^−/−^ mice also accrue lysosomal proteins such as Lamp1 and cathepsin-L, as well as ATP5G – a mitochondrial protein characteristically stored in lysosomes of CSPα loss-of-function models and ANCL patient neurons (**Supplementary Fig. S1**). The enhanced accumulation of lysosomal contents in Tg-αSyn^A53T^/CSPα^−/−^ brains, together with the exaggerated buildup of αSyn aggregates in these brains gave us the clue that αSyn aggregates may be depositing in the lysosomes.

We next aimed to determine whether αSyn aggregates accumulate within lysosomes, even in the absence of CSPα^−/−^-driven lysosomal pathology. Thus, we tested whether aged Tg-αSyn^A53T^ mice, which display severe αSyn-driven pathology, have αSyn aggregates within their lysosomes.

We found pathogenic αSyn species, but not monomeric αSyn, in dextranosomes (heavy lysosomes loaded with dextran to enhance purity) isolated from the brains of 6 month old Tg-αSyn^A53T^ mice (**Fig. 1 a-c**). These lysosomal fractions precisely coincided with pathogenic αSyn species, and had highly enriched lysosomal proteins as well as cathepsin-D activity, with negligible contamination from other organelle protein-markers or mitochondrial and peroxisomal enzyme activities (**Fig. 1 a-c**). This experiment indicates that isolated intact lysosomes from aged Tg-αSyn^A53T^ mice contain pathogenic αSyn species.

**Figure 1.**
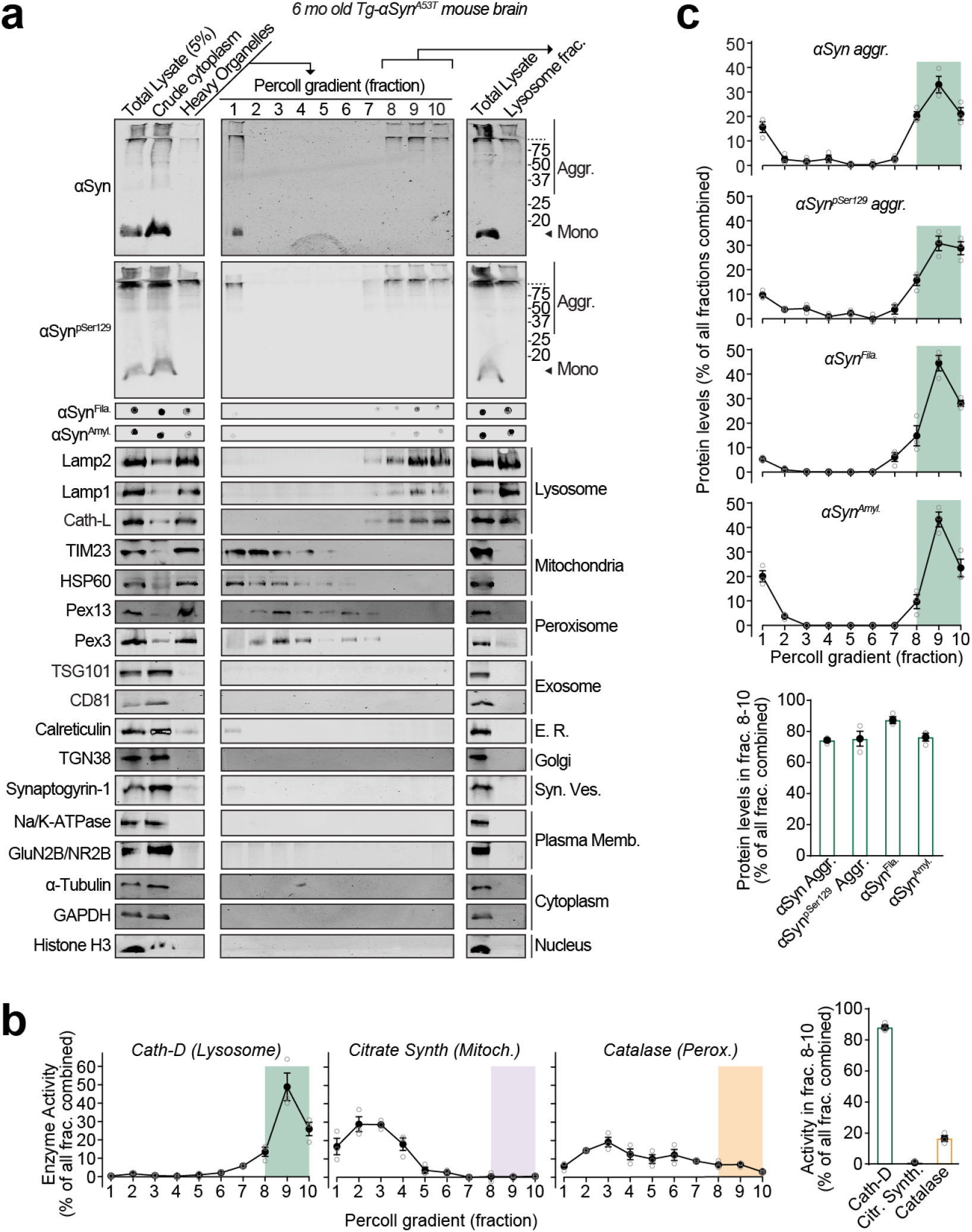
Pathogenic αSyn aggregates accumulate within lysosomes of aged Tg-αSyn^A53T^ mice. Lysosomes were isolated from 6 month old Tg-αSyn^A53T^ mouse brains via Percoll gradient centrifugation of the heavy organelle fraction – which contained peroxisomes, heavy lysosomes loaded *in vivo* with dextran-70, and mitochondria swollen *ex vivo* by CaCl_2_. (**a**) Lysosome (dextranosome) enrichment was determined by immunoblotting for markers of indicated organelles, compared to the respective levels in the total lysate input. (**b**) Left panels – Activities of enzymes contained within lysosomes (cathepsin-D), mitochondria (citrate synthase), and peroxisomes (catalase) were measured, testing for isolation of intact organelles. Right panel – Summary graph of enzyme activity present in the combined “lysosomal fractions” (fractions 8-10). (**c**) Top panels – Levels in each Percoll gradient fraction of pathogenic αSyn species: aggre-gated (αSyn Aggr), aggregates phosphorylated at Ser129 (αSyn^pSer129^ Aggr), amyloid-type (αSyn^Amyl^), and filamentous (αSyn^Fila^). Bottom panel – Summary graph of these αSyn species present in the combined “lysosomal fractions” (fractions 8-10). (n=3). (**b-c**) All data in are shown as means ± SEM, where ‘n’ represents mouse brains.

### Pathogenic species of αSyn accumulate within the lumen of neuronal lysosomes

The lysosomes studied above are a mixture of lysosomes from all brain cells. To isolate and study lysosomes specifically from neurons, we generated transgenic mice expressing ^HA^Lamp1^Myc^ – lysosomal protein Lamp1 epitope-tagged on its luminal domain with tandem 2xHA, and on its cytoplasmic tail with 2xmyc – driven by the neuron-specific synapsin-I promoter (**Supplementary Fig. S2 a-d**), enabling affinity-isolation of neuronal lysosomes from mouse brains. The Tg-^HA^Lamp1^Myc^ mice were then crossed to the Tg-αSyn^A53T^ mice to generate Tg-αSyn^A53T^/Tg-^HA^Lamp1^Myc^ progeny (**Supplementary Fig. S3 a-g**). Immuno-isolation experiments from aged Tg-αSyn^A53T^/Tg-^HA^Lamp1^Myc^ mouse brains indicated that pathogenic versions of αSyn co-isolate with neuronal lysosomes (**Fig 2a**). We found that ~10% of the SDS-resistant αSyn aggregates, and ~20% of the more mature αSyn aggregates (filamentous and amyloid-type) reside within neuronal lysosomes (**Fig. 2a**). This result indicates that a non-trivial portion of pathogenic αSyn species in the brain are found in neuronal lysosomes.

**Figure 2.**
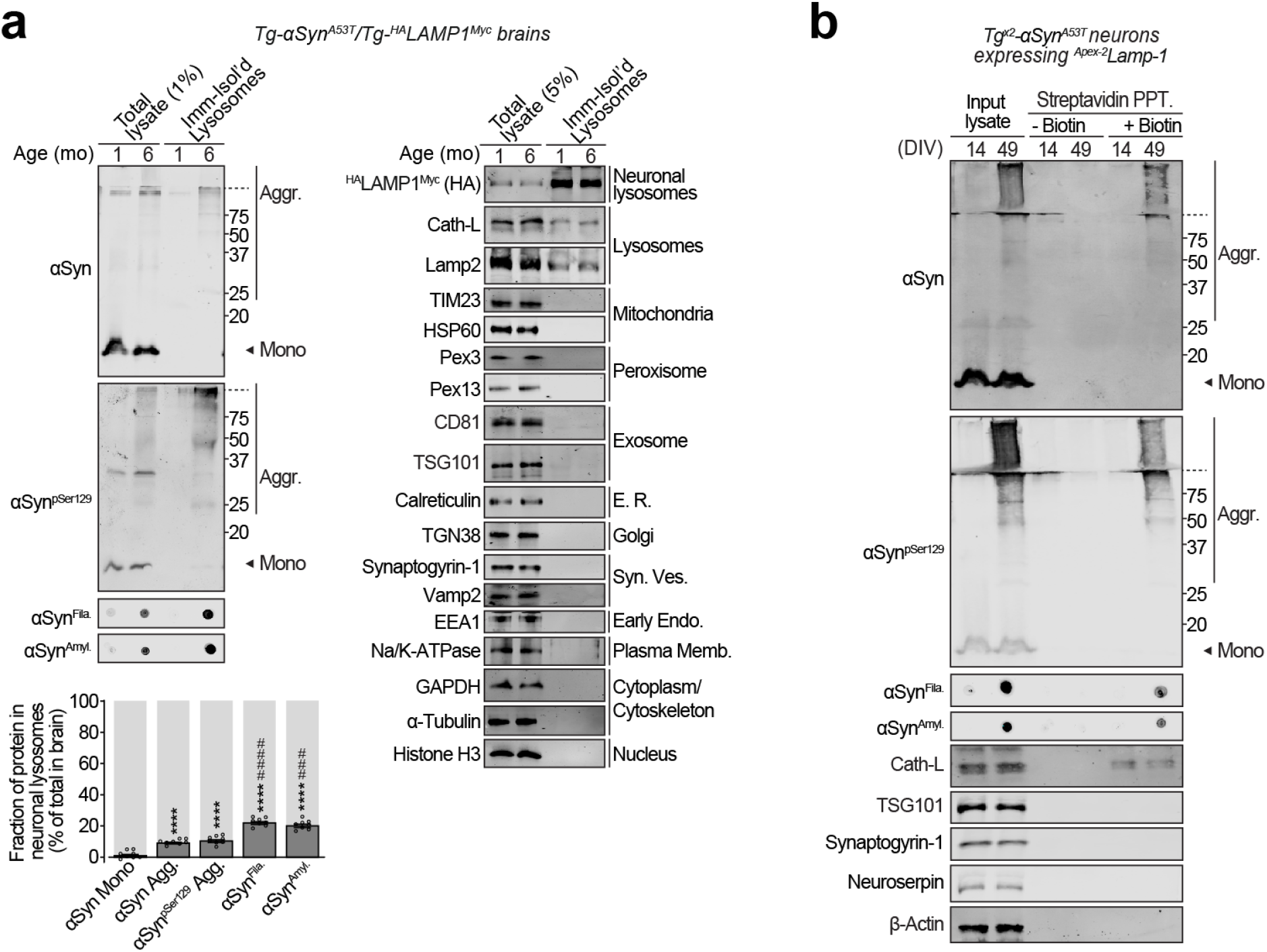
Pathogenic αSyn species accumulate within neuronal lysosomes in mouse brains and primary neurons. (**a**) Neuronal lysosomes immunoisolated from the brains of 1 and 6 month old Tg-αSyn^A53T^/^HA^Lamp1^Myc^ mice using antimyc antibody were analyzed by immunoblotting for levels of αSyn, αSyn^pSer129^, α-Syn^Fila^, and αSyn^Amyl^ (Mono = monomer, Aggr = aggregates). Fraction of these αSyn species residing within the lysosomes at 6 months of age (bottom graph); this was back-calculated from the fraction of each αSyn species immuno-captured, normalized to the fraction of neuronal lysosomes captured – indicated by the fraction of ^HA^Lamp1^Myc^ captured. Immunoblots against markers of lysosomes and of other organelles are also shown (n=8). (**b**) Tg^x2^-αSyn^A53T^ primary cultures lentivirally expressing a synapsin-1 promoter-driven ^Apex-2^Lamp1 construct were subjected to proximity-labeling with biotin at 14 or 49 days *in vitro* (DIV). Labeled proteins were precipitated on streptavidin-coated magnetic beads and immunoblotted for the indicated pathogenic αSyn species, as well as for marker proteins cathepsin-L (lysosomes), TSG101 (exosomes), synaptogyrin-1 (synaptic vesicles), neuroserpin (constitutively secreted neuronal protein), and β-actin (cytosol) (representative of n=3). Data represent means ± SEM. Each ‘n’ corresponds to independently aged mouse littermates in (**a**) and independent primary neuron cultures in (**b**). ****P<0.0001 by RM 1-way ANOVA with Dunnett multiple-comparison correction; and ^###^P<0.001; ^####^P<0.0001 by non-parametric Friedman test with Dunn’s multiple-comparison adjustment.

To understand the fate of these lysosomal αSyn aggregates, we developed a neuronal model which accumulates pathogenic αSyn species. Compared to Tg-αSyn^A53T^ mice, homozygous Tg-αSyn^A53T^ mice (Tg^x2^-αSyn^A53T^), which express twice the amount of transgenic αSyn^A53T^, have shorter lifespans, earlier onset of neuromuscular pathology, and begin accumulating pathogenic αSyn species earlier – by only 6 weeks of age (**Suppl. Fig. S4 a-f**) – which is achievable in long-lived primary neuron cultures. Accordingly, primary neurons from Tg^x2^-αSyn^A53T^ mice also accrued pathogenic αSyn species by 6 weeks, including aggregates detectable by antibodies against αSyn fibrils and amyloid-type conformations (**Suppl Fig. S4g**).

To specifically label and track the pathogenic αSyn species located within neuronal lysosomes, we targeted Apex-2 to the lysosomal lumens of Tg^x2^-αSyn^A53T^ primary neurons. We lentivirally expressed a synapsin-1 promoter-driven construct, comprised of Apex-2 fused to the luminal domain of a truncated version of Lamp1 (^Apex-2^Lamp1) (**Suppl Fig. S5 a-c**), which proximity-labels the luminal contents of neuronal lysosomes with biotin. In these Tg^x2^-αSyn^A53T^ neurons expressing ^Apex-2^Lamp1, we found that pathogenic αSyn species as well as lysosomal proteins (typified by cathepsin-L) were biotinylated (**Fig. 2b**), confirming that pathogenic αSyn species accumulate within the lumen of neuronal lysosomes.

### Pathogenic αSyn species are secreted from neurons via SNARE-dependent lysosomal exocytosis

Release of αSyn from neurons, particularly the seeding-competent pathogenic versions of αSyn, is considered a key step in the spatial progression of synucleinopathies (Peng et al., 2020), and αSyn has been documented in extracellular fluids of human PD patients (El-Agnaf et al., 2003). We also found pathogenic species of αSyn in the cerebrospinal fluid of aged Tg-αSyn^A53T^ mice (**Suppl. Fig. S6**). Importantly, this extracellular pool of αSyn species appears not to be enveloped in membranes, as they were not protected from proteinase K digestion (**Suppl. Fig. S6**). Proteolytic susceptibility was also unaffected by the presence of detergent, akin to the lysosomal luminal protein cathepsin-L and the constitutively secreted neuronal protein neuroserpin (**Suppl. Fig. S6**). This was contrary to the well-protected contents of exosomes, typified by TSG101, which becomes susceptible to proteolysis only in the presence of detergent (**Suppl. Fig. S6**). Accumulation inside lysosomes of the pathogenic αSyn species, together with their un-enveloped presence in the extracellular space, prompted us to investigate whether lysosomes are releasing αSyn aggregates contained inside them.

We thus collected the medium from Tg^x2^-αSyn^A53T^ primary neurons, in which the lysosomal contents had been biotinylated via neuron-specific expression of ^Apex-2^Lamp1 (as in **Fig. 2b** and **Suppl. Fig. S5**). We found that lysosomal contents, typified by cathepsin-L, as well as pathogenic αSyn species could be precipitated from the medium via their biotin-modification (**Fig. 3a**). In contrast, exosomal contents, typified by TSG101, were not biotinylated/precipitated. This result points to lysosomal exocytosis – SNARE-dependent fusion of lysosomes with the plasma membrane – as the likely mechanism for the exit of αSyn aggregates from neurons.

**Figure 3.**
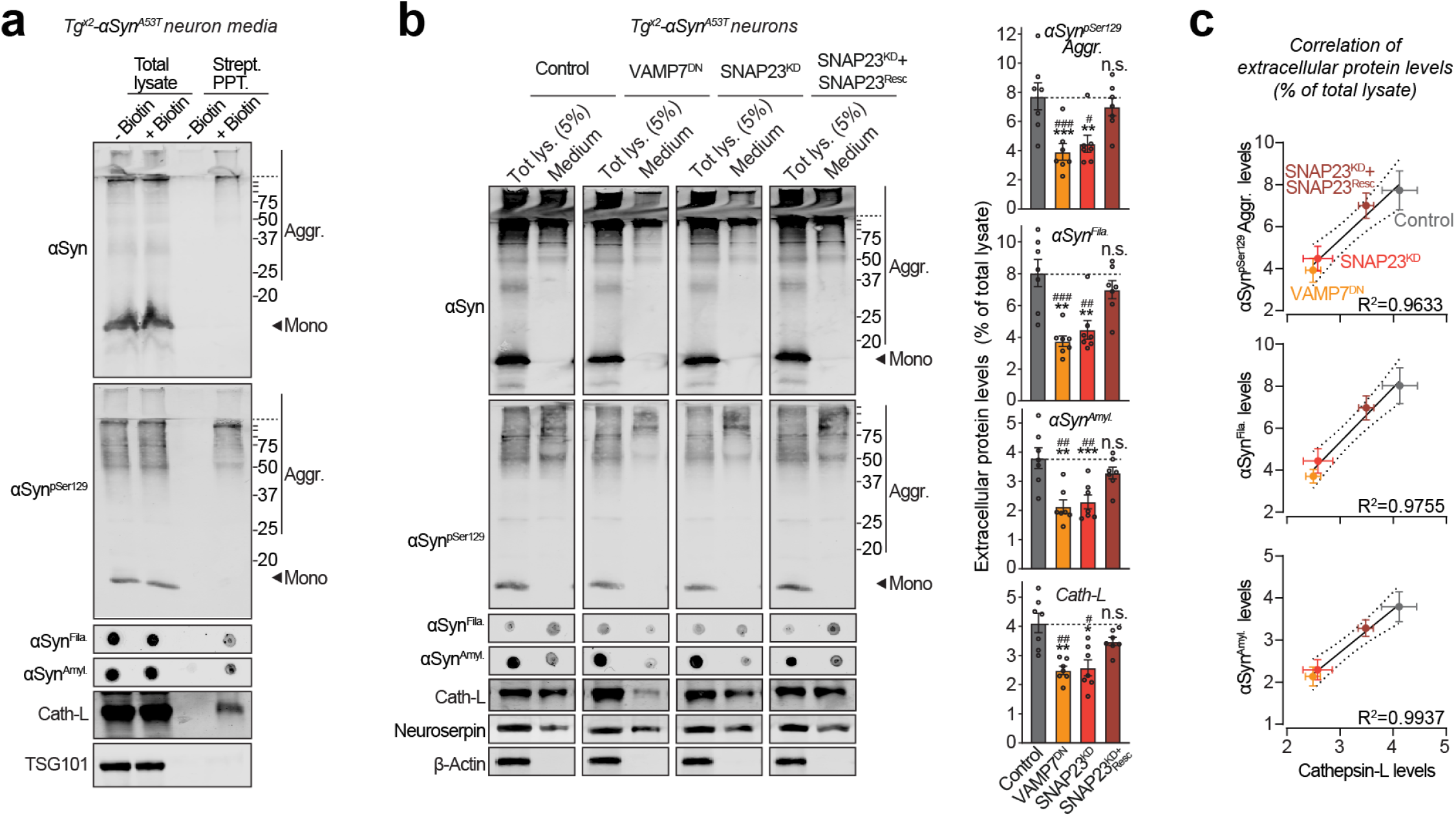
Pathogenic αSyn species are secreted from neurons via SNARE-dependent lysosomal exocytosis. (**a**) Tg^x2^-αSyn^A53T^ primary neurons lentivirally expressing a synapsin-1 promoter-driven ^Apex-2^Lamp1 (infected on DIV 7) were subjected to proximity-labeling on DIV 47. Biotinylated proteins released into the media during 48 h (by DIV 49) were precipitated on streptavidin-magnetic beads and immunoblotted for the indicated αSyn species (αSyn, αSyn^pSer129^, αSyn^Amyl^, and αSyn^Fila^), lysosome luminal protein cathepsin-L, and exosome luminal protein TSG101. Strept. PPT. = strep-tavidin precipitate. (representative of n=3). (**b**) Media collected over 2 days (DIV 47-49) from Tg^x2^-αSyn^A53T^ primary neurons lentivirally expressing control (GFP), GFP-VAMP7^DN^ fragment, SNAP23^KD^ shRNA, and SNAP23^KD^ shRNA plus knockdown-resistant SNAP23 rescue-construct (infected on DIV 7), were immunoblotted for the indicated αSyn species (αSyn, αSyn^pSer129^, αSyn^Amyl^, and αSyn^Fila^), cathepsin-L (lysosome luminal), neuroserpin (constitutively secreted from neurons), and β-actin (cytoplasmic). For quantification (graphs on the right) protein level in medium was normalized to its levels in total cellular lysate (5% loaded) (n=7). (**c**) Correlation between the levels of pathogenic αSyn species and the levels of cathepsin-L released in the medium, upon the indicated manipulation of SNARE-dependent lysosomal exocytosis; derived from data shown above in panel (b). Linear regression is shown with dotted lines depicting the 95% confidence intervals, and Pearson’s correlation coefficient is indicated on the bottom right of each graph. (n=7). All data represent means ± SEM. Each ‘n’ corresponds to a separate mouse litter used for a batch of neuronal culture and infection. In (**b**) *P<0.05; **P<0.01; ***P<0.001; by RM 1-way ANOVA with Dunnett multiple-comparison correction; and ^#^P<0.05; ^##^P<0.01; ^###^P<0.001 by non-parametric Friedman test with Dunn’s multiple-comparison adjustment.

To test this hypothesis, we modulated the SNARE proteins responsible for lysosome-to-plasma membrane fusion. First, we found VAMP7 as the prominent v-SNARE in immunoisolated lysosomes (**Suppl. Fig. S7a**). We then screened for the Qb-type t-SNAREs which interact and thus co-immunoprecipitate with VAMP7. SNAP23 was the clearest Qb/t-SNARE candidate (**Suppl. Fig. S7a**). Based on these results and prior studies validating them (Rao et al., 2004), we tested four strategies to disrupt lysosomal exocytosis: shRNA knockdown of either SNAP23 (SNAP23^KD^) or VAMP7 (VAMP7^KD^), and overexpression of dominant negative fragments of either SNAP23 (SNAP23^DN^) or VAMP7 (VAMP7^DN^). In wild type primary neurons, the release of cathepsin-L was used to signal lysosomal exocytosis, opposed to the constitutively secreted neuroserpin. VAMP7^KD^ and SNAP23^DN^ had no significant effect on lysosomal exocytosis, but VAMP7^DN^ severely reduced it by ~80% and SNAP23^KD^ partially reduced it by ~40% (**Suppl. Fig. S7 b-c**). Therefore, we applied the latter two strategies to disrupt lysosomal exocytosis next.

Importantly, in Tg^x2^-αSyn^A53T^ neurons, both VAMP7^DN^ and SNAP23^KD^ strategies reduced the release of pathogenic αSyn species into the medium, and the effect of SNAP23^KD^ was rescued by overexpression of a knockdown-resistant version of wild type SNAP23 (**Fig. 3b**). Moreover, the release of pathogenic αSyn species was significantly correlated with the change in lysosomal exocytosis, measured as cathepsin-L released into the medium (**Fig. 3 b-c**). These results suggest that SNARE-dependent exocytosis of lysosomes is essential and rate-determining for the release of pathogenic αSyn species from neurons.

### αSyn species released via lysosomal exocytosis can seed aggregation of purified recombinant αSyn

Pathogenic spread requires that the released αSyn is able to template or “seed” the assembly of amyloid-type aggregates from monomeric αSyn. To test whether the pathogenic αSyn species exocytosed from neurons are seeding-competent, we performed *in vitro* aggregation of purified recombinant myc-tagged wild type αSyn (**Suppl. Fig. S8**), in the presence of extracellular media collected from neuron cultures generated either from wild type mice, or from Tg^x2^-αSyn^A53T^ mice with or without lentiviral expression of VAMP7^DN^ fragment (**Fig. 4 a-c**). Aggregation kinetics, measured by amyloid-binding fluorescent dye K114 (**Fig. 4a**) or Thioflavin-T (**Fig. 4b**), were enhanced by medium from Tg^x2^-αSyn^A53T^ neurons, compared to the medium from wild type neurons. Suppressing lysosomal exocytosis in the Tg^x2^-αSyn^A53T^ neurons, via VAMP7^DN^ expression, diminished the aggregation-promoting effect of the Tg^x2^-αSyn^A53T^ medium (**Fig. 4 a-b**). When myc-αSyn aggregation kinetics were measured by quantitative immunoblotting for either disappearance of αSyn monomers or for the appearance of filamentous and amyloid-type aggregates, we again found that compared to the wild type neuron medium, aggregation was augmented by the Tg^x2^-αSyn^A53T^ medium, and the expression of VAMP7^DN^ fragment diminished this effect (**Fig. 4c**). In all the assays, a lag-time of nearly 2 weeks is apparent before the appearance of amyloid signal with the wild type medium (**Fig. 4a-c**). Importantly, this lag is shortened or eliminated by the aggregates in Tg^x2^-αSyn^A53T^ medium (**Fig. 4a-c**), with a reduced effect when lysosomal exocytosis is suppressed by VAMP7^DN^ fragment (**Fig. 4a-c**). This seeding activity of the lysosomally released αSyn species indicates that they are capable of amyloid-nucleation via permissive templating.

**Figure 4.**
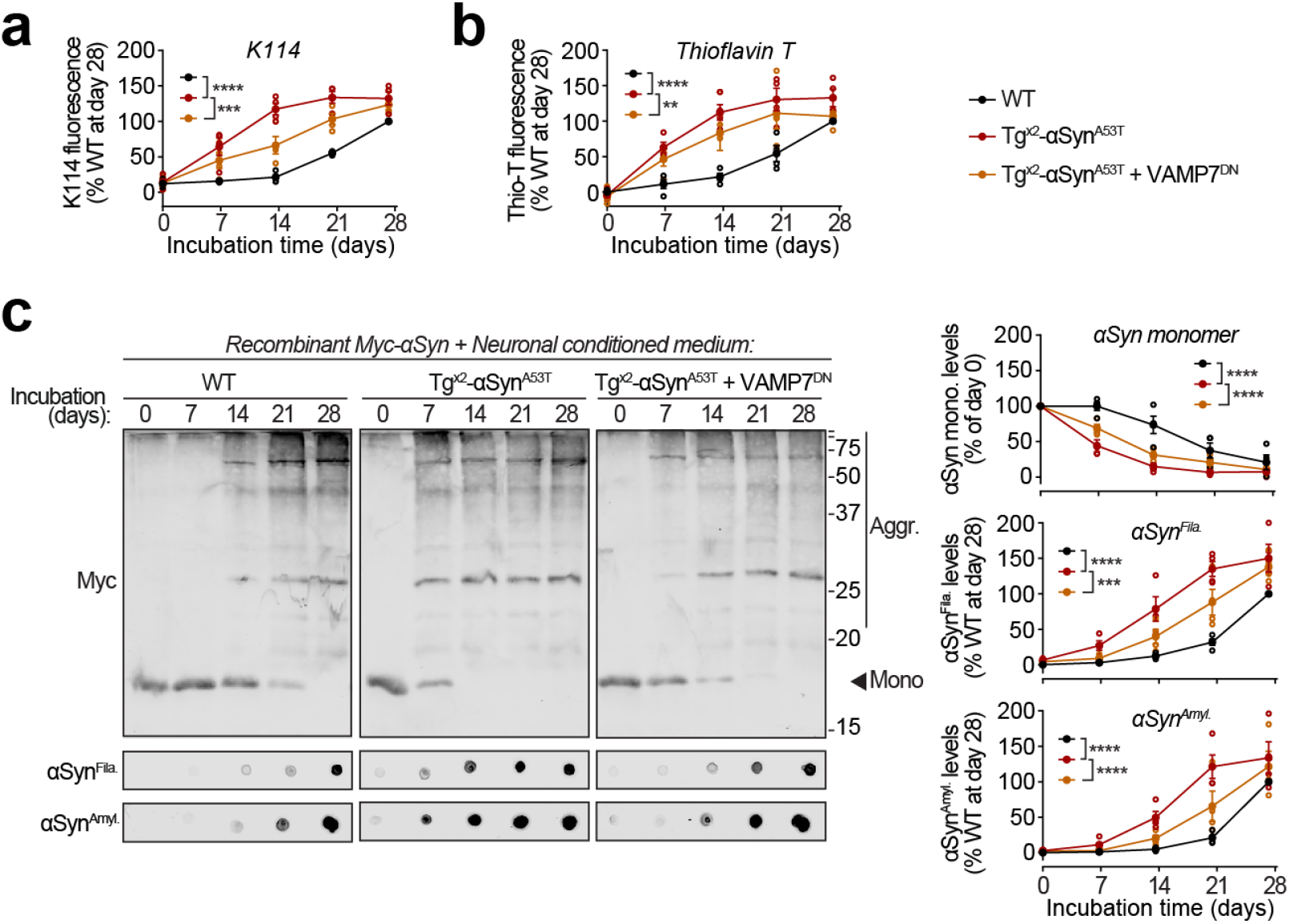
Seeding of recombinant αSyn aggregation by pathogenic αSyn species exocytosed from neurons. (**a-c**) Recombinant purified myc-αSyn was shaken at 37°C in presence of concentrated extracellular medium from mouse cortical neuron cultures collected over 7 days (DIV 42-49), generated either from wild type (WT) mice, or from Tg^x2^-αSyn^A53T^ mice with or without lentiviral expression of VAMP7 dominant-negative fragment (VAMP7^DN^; infected at DIV 7). Aggregation of myc-αSyn was analyzed at the indicated days of incubation by the following assays: (**a**) Congo-red derivative, amyloid-binding dye K114 fluorescence at 390/535 nm (n=4). (**b**) Amyloid-binding dye Thioflavin-T fluorescence at 450/485 nm (n=4). (**c**) Quantitative immunoblotting for the myc epitope-tag, where aggregation is measured as disappearance of monomeric myc-αSyn (top; n=4); dot-blotting for filamentous myc-αSyn aggregates using αSyn^Fila^ antibody (middle; n=4); and dot-blotting for amyloid-type myc-αSyn aggregates using αSyn^Amyl^ A11 antibody (bottom; n=4). All data represent means ± SEM, where each ‘n’ is an independent aggregation experiment. *P<0.05; **P<0.01; ***P<0.001; ****P<0.0001 by RM 2-way ANOVA.

## DISCUSSION

Release of αSyn aggregates from neurons is thought to transmit neuropathology to other yet-unaffected neurons. Here we find that pathogenic αSyn species accumulate within neuronal lysosomes in mouse brains and in primary neurons, and are then released from neurons via SNARE-dependent lysosomal exocytosis.

It is important to note that for αSyn pathology to transmit from neuron-to-neuron, the lysosomal exocytosis mechanism is more in line with the following observations, which underline that the extracellular pool of pathogenic αSyn is likely not membrane-enveloped as in exosomes, extracellular vesicles, or nanotubes, but is non-enveloped: a) the stark pathology and rapid propagation caused by inoculated pre-formed fibrils (Luk et al., 2012; Luk et al., 2009; Paumier et al., 2015; Rey et al., 2018; Rey et al., 2013; Volpicelli-Daley et al., 2011b), b) the efficacy of humoral and cellular immunotherapy against αSyn (reviewed in (Chatterjee and Kordower, 2019)), c) the efficacy of passive immunization using antibodies against αSyn epitopes (reviewed in (Chatterjee and Kordower, 2019)), and d) much of the extracellular αSyn has been found as non-enveloped (Borghi et al., 2000; El-Agnaf et al., 2003; Sung et al., 2005), while a minority of αSyn is contained in extracellular vesicles or exosomes (Ejlerskov et al., 2013; Emmanouilidou et al., 2010; Jang et al., 2010).

Key gaps still remain in how the pathogenic αSyn species are targeted and trafficked to lysosomes, as well as in the fate of extra cellular αSyn aggregates following their release from neurons:

The aggregates/fibrils of αSyn could form inside the acidified lumen of lysosomes (Buell et al., 2014), either from smaller oligomers or monomers, especially if their degradation is delayed due to reduced/disrupted lysosomal activity (Malik et al., 2019). Yet, we found no presence of αSyn monomers within isolated lysosomes, making this possibility less likely. In contrast, macro-autophagy of ready-formed aggregates from the cytoplasm, termed “aggrephagy”, is a more likely mechanism, following from the well-recorded ability of autophagosomes to encapsulate large cytoplasmic structures including aggregates (Yamamoto and Simonsen, 2011), and target them to lysosomes.

Once released from the neurons, the fate of extracellular/interstitial αSyn aggregates also remains unclear, and can follow multiple possible scenarios: uptake into microglia (Lee et al., 2008), astrocytes (Loria et al., 2017), and/or oligodendrocytes (Reyes et al., 2014), and/or drainage via glymphatics (Rasmussen et al., 2018; Smith and Verkman, 2018) or blood-flow (El-Agnaf et al., 2003; Foulds et al., 2013). However, for the extracellular αSyn aggregates to cause the stereotypic “spread” of pathology via a prion-like etiology, other neurons will have to take up the released aggregates, and this process remains even less understood than the glial uptake or clearance scenarios.

The results presented here provide evidence of the accumulation of pathogenic αSyn species in neuronal lysosomes, and pinpoint to lysosomal exocytosis as a pathway of release for these pathogenic αSyn species into the extracellular milieu. This pathway, at first sight, appears to have relevance to the release of undegraded aggregates composed of other proteins as well that transmit pathology from cell-to-cell (reviewed in (Peng et al., 2020)), and follow-up studies will be needed to test this speculation.

---

**Supplementary Figure S1.**
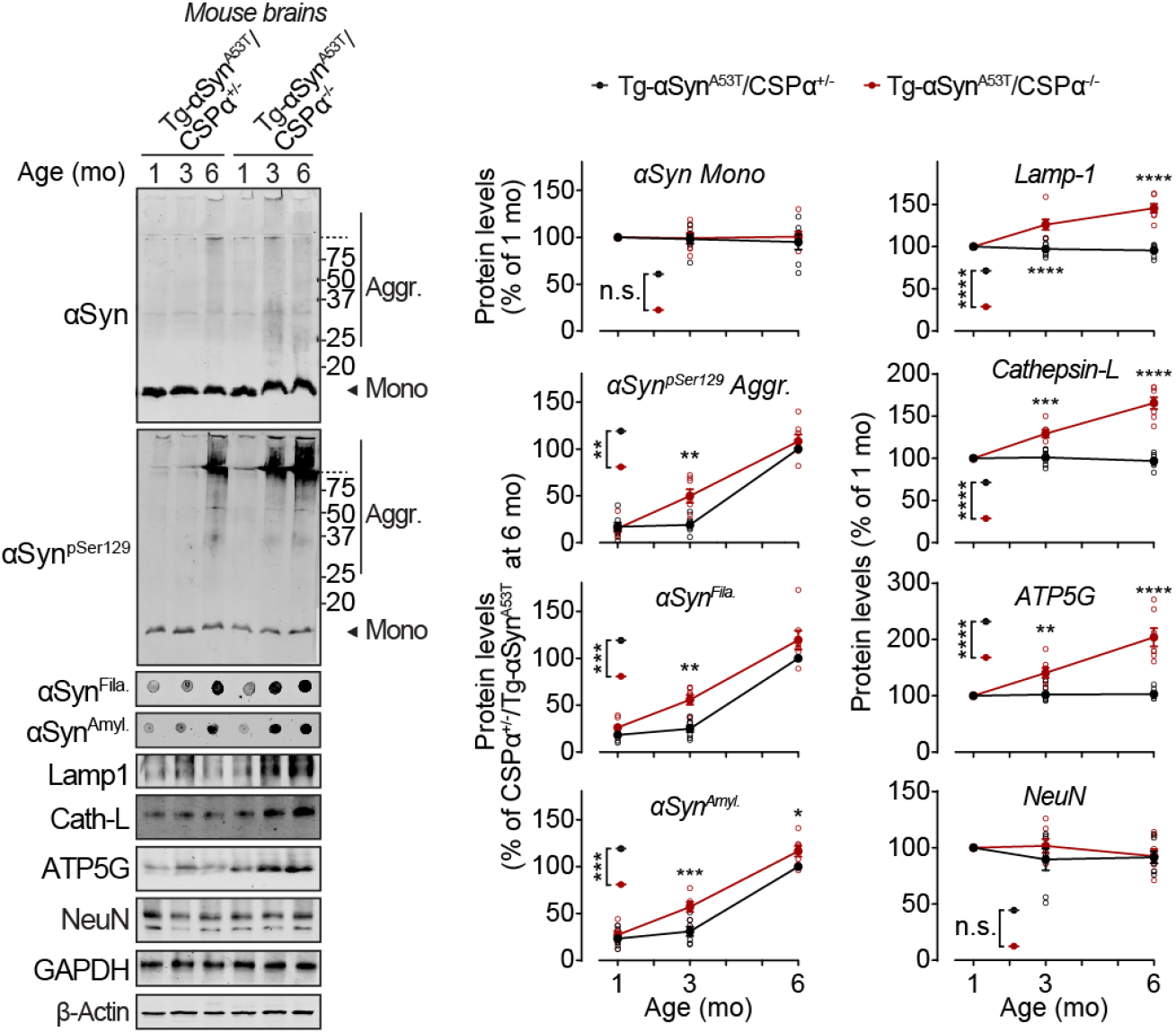
Accelerated accumulation of pathogenic αSyn aggregates and lysosomal proteins in Tg-αSyn^A53T^ mouse brains due to the loss of CSPα function. Brains collected from Tg-αSyn^A53T^/CSPα^+/−^ and Tg-αSyn^A53T^/CSPα^−/−^ littermates at 1, 3 and 6 months of age were analyzed by quantitative immunoblotting for the following versions of αSyn: monomeric (αSyn Mono), phosphorylated at Ser129 (αSyn^pSer129^), filamentous (αSyn^Fila^), and amyloid-type (αSyn^Amyl^); for lysosomal proteins Lamp1 and cathepsin-L (Cath-L); for ATP5G, which accumulates in lysosomes storage caused by loss of CSPα function; as well as for neuronal marker NeuN. These were all normalized to β-actin levels. Mono = monomer; Aggr. = aggregates. (n=7) All data are shown as means ± SEM, where ‘n’ represents littermate mouse brains. *P<0.05; **P<0.01; ***P<0.001; ****P<0.0001 by RM 2-way ANOVA with Bonferroni multiple comparisons post-test.

**Supplementary Figure S2.**
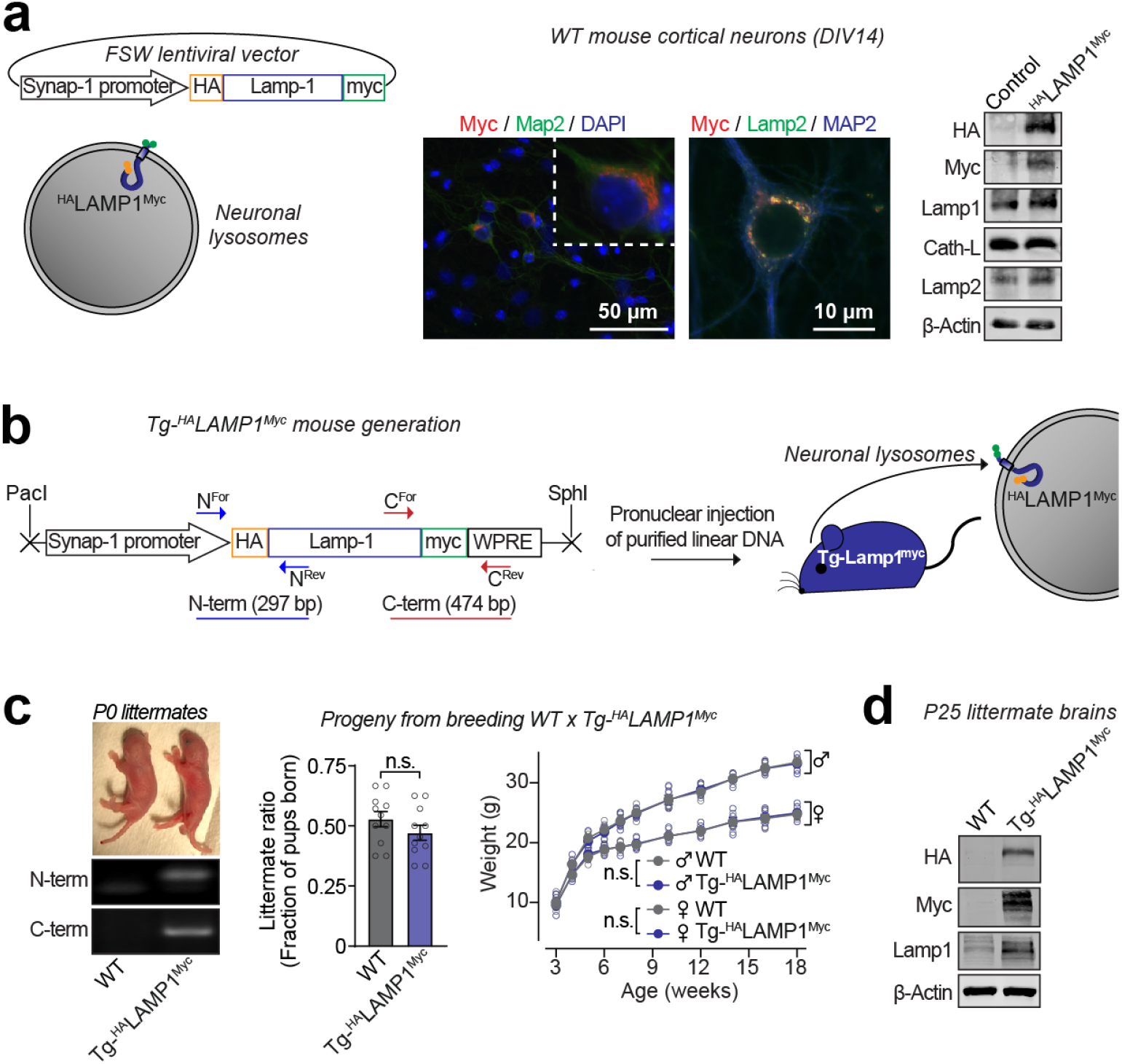
Development of a transgenic mouse model to isolate neuronal lysosomes from brains. (**a**) ^HA^Lamp1^Myc^ construct lentivirally expressed via neuron-specific synapsin-1 promoter (schematics on left), localizes in neurons marked by Map-2 (and not in surrounding glia seen in DAPI panel), and co-localizes with the lysosomal protein Lamp2 in primary neuron cultures (immunofluorescence in middle panels). Expression of full length protein ^HA^Lamp1^Myc^ is confirmed by immunoblotting for its epitope tags: N-terminal HA tag and C-terminal myc tag. (**b**) This lentiviral vector was used to generate Tg-^HA^Lamp1^Myc^ mice via pronuclear injection of linearized DNA comprising the synapsin-1 promoter and the ^HA^Lamp1^Myc^ cDNA. (**c**) Tg-^HA^Lamp1^Myc^ mice were PCR-genotyped using primer-pairs at the N- and the C-terminal ends of the ^HA^Lamp1^Myc^ cDNA. Single insertion-site for the transgene was suggested by equal numbers of WT and Tg-^HA^Lamp1^Myc^ pups born per litter (n=11 litters). There was also no effect of the transgene on birth-ratio and growth (indicated by weight gain; n=5 per group) of the Tg-^HA^Lamp1^Myc^ mice compared to WT mice. (**d**) Immunoblots show that ^HA^Lamp1^Myc^ protein is detectable in mouse brains by immunoblots against the N-terminal HA tag and the C-terminal myc tag. All data represent means ± SEM. n.s. = not significant, by 2-tailed Student’s t-test for littermate ratio and by RM 2-way ANOVA for weight gain over age. *Note:* Further characterization of the Tg-^HA^Lamp1^Myc^ mice is included in **Supplementary Fig. S3**.

**Supplementary Figure S3.**
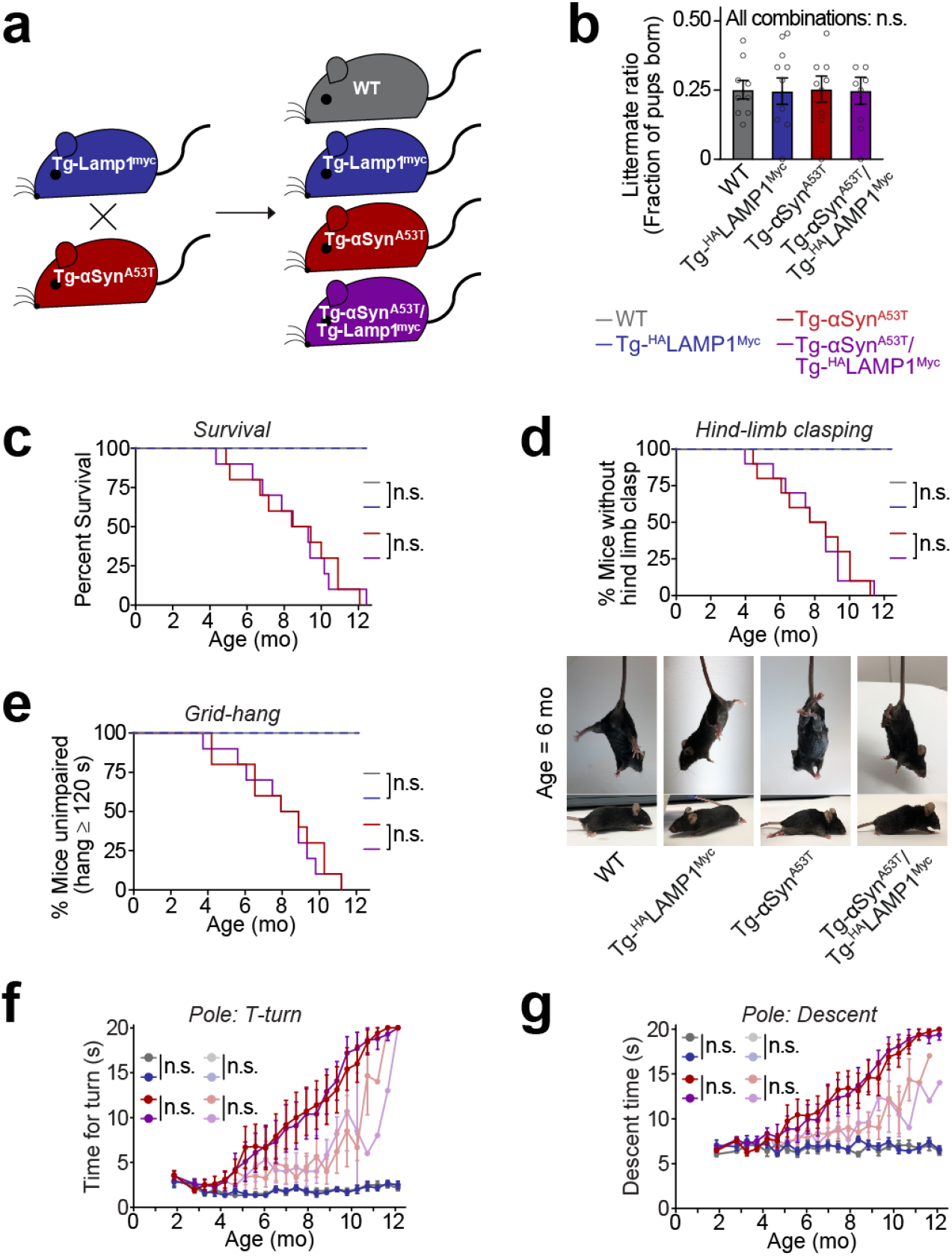
In Tg-αSyn^A53T^/Tg-^HA^Lamp1^Myc^ double-transgenic mice, the neuromuscular impairments driven by αSyn^A53T^ transgene remain unaffected by the ^HA^Lamp1^Myc^ transgene. (**a**) Tg-αSyn^A53T^ mice were crossed to Tg-^HA^Lamp1^Myc^ mice to generate the Tg-αSyn^A53T^/Tg-^HA^Lamp1^Myc^ progeny. (**b**) Equal numbers of pups with the four resultant genotypes (WT, Tg-^HA^Lamp1^Myc^, Tg-αSyn^A53T^, and Tg-αSyn^A53T^/Tg-^HA^Lamp1^Myc^) were born from crossing Tg-αSyn^A53T^ and Tg-^HA^Lamp1^Myc^ parents (n=9 litters). Tg-αSyn^A53T^/Tg-^HA^Lamp1^Myc^ and Tg-αSyn^A53T^ mice were indistinguishable in (**c**) lifespan, as well as in the onset and course of neuromuscular deterioration (**d-g**): Signs of progressive neuromuscular debility preceded death, as measured by (**d**) onset of hind-limb clasping, (**e**) grid-hang (limb strength), (**f**) T-turn on pole-test (bradykinesia), and (**g**) descent on pole test (motor coordination) (n=10 mice per group). (**f-g**) Mice that died were scored as 20 s (maximum measurement) for the rest of the trial – not as an actual measurement, but as a place holder. The same data are shown in the lighter shaded graphs without the 20 s placeholder, as dead animals are eliminated from the cohort over time. WT and Tg-^HA^Lamp1^Myc^ mice showed no impairments in these measurements (n=5 mice per group). (**b and f-g**) Data represent means ± SEM. n.s. = not significant, by RM 1-way ANOVA in (**b**), Log-rank (Mantel-Cox) test for comparisons of Kaplan-Meier curves (**c-e**), RM 2-way ANOVA (dark shaded graphs) and mixed-effects analysis (lighter shaded graphs) for pole turn and pole descent test (**f-g**).

**Supplementary Figure S4.**
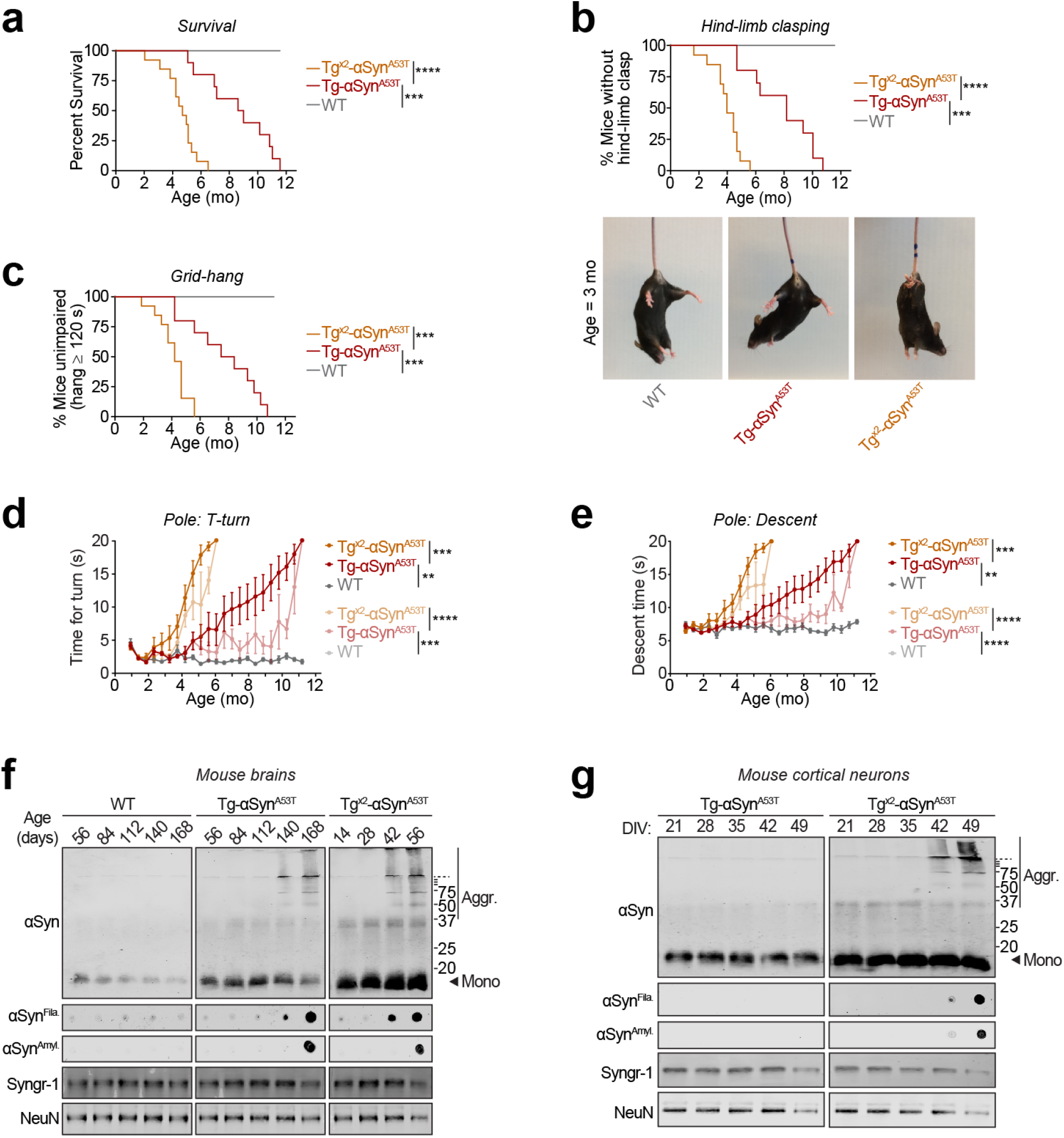
Characterization of the homozygous Tg^x2^-αSyn^A53T^ mice and primary neurons. (**a**) Doubling the dose of αSyn^A53T^ in Tg^x2^-αSyn^A53T^ mice resulted in accelerated mortality, nearly halving the lifespan of Tg-αSyn^A53T^ mice. This is accompanied by accelerated onset of neuromuscular impairment in Tg^x2^-αSyn^A53T^ mice, as measured by (**b**) onset of hind-limb clasping, (**c**) onset of grid-hang impairment (limb strength) (**d**) T-turn on pole-test (bradykinesia), and (**e**) descent on pole test (motor coordination). WT mice showed no impairments during the time course (n=10 Tg-αSyn^A53T^; n=13 Tg^x2^-αSyn^A53T^; n=6 WT mice). (**d-e**) Mice that died were scored as 20 s (maximum measurement) for the rest of the trial – not as an actual measurement, but as a place holder. The same data are shown in the lighter shaded graphs without the 20 s placeholder, as dead animals are eliminated from the cohort over time. (**f**) In addition to increased levels of αSyn monomer, αSyn aggregates are detected in Tg^x2^-αSyn^A53T^ brains much earlier than in Tg-αSyn^A53T^ brains. (**g**) Tg^x2^-αSyn^A53T^ primary neuron cultures recapitulate the accelerated accumulation of pathogenic αSyn species. (**a-c**) ***P<0.001 and ****P<0.0001 by Log-rank (Mantel-Cox) test. (**d-e**) Data represent means ± SEM, **P<0.01; ***P<0.001; ****P<0.0001 by RM 2-way ANOVA (dark shagged graphs) and mixed-effects analysis (lighter shaded graphs); analysis between Tg^x2^-αSyn^A53T^ and Tg-αSyn^A53T^ is up to 6 mo.

**Supplementary Figure S5.**
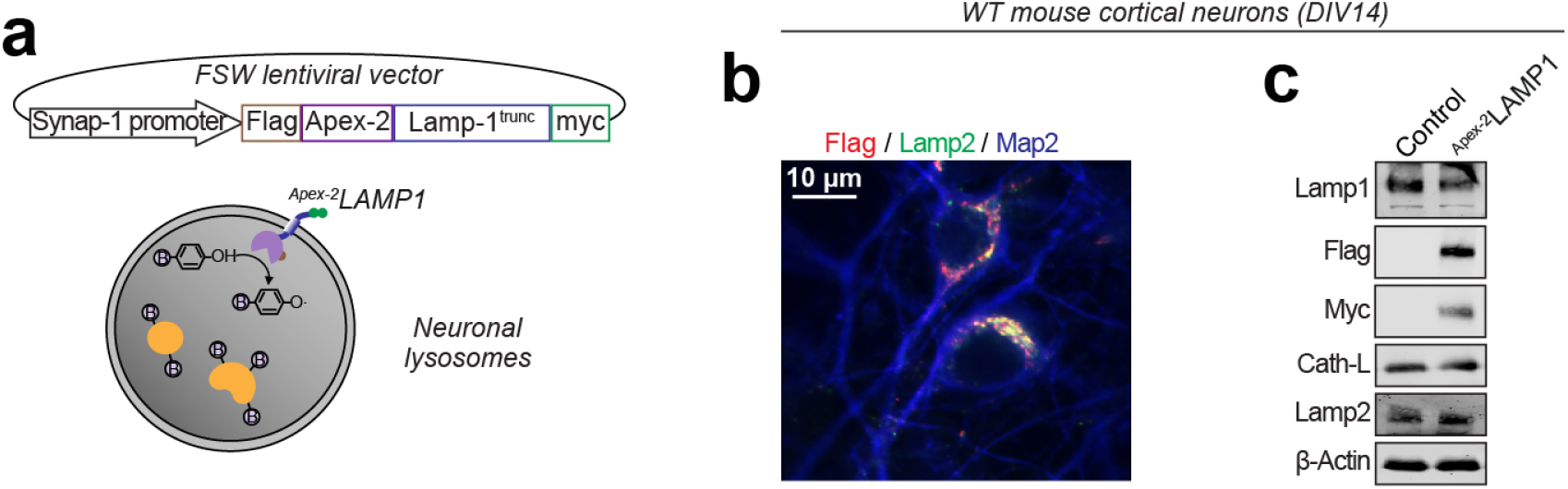
Generation of ^Apex-2^Lamp1 chimeric protein for labeling lysosomal luminal proteins in primary neurons. (**a**) Apex-2 was fused to the N-terminus of truncated Lamp1, and expressed via neuron-specific synapsin-1 promoter, allowing it to biotinylate lysosomal luminal proteins. (**b**) Lentivirally expressed ^Apex-2^Lamp1 (infected on DIV 7, immunostained on DIV14) co-localizes with lysosomal protein Lamp2 in primary neurons. Neuronal somal and dendrites are marked by Map2. (**c**) Expression of ^Apex-2^Lamp1 in primary neurons is confirmed by immunoblotting for its epitope tags: N-terminal FLAG and C-terminal myc.

**Supplementary Figure S6.**
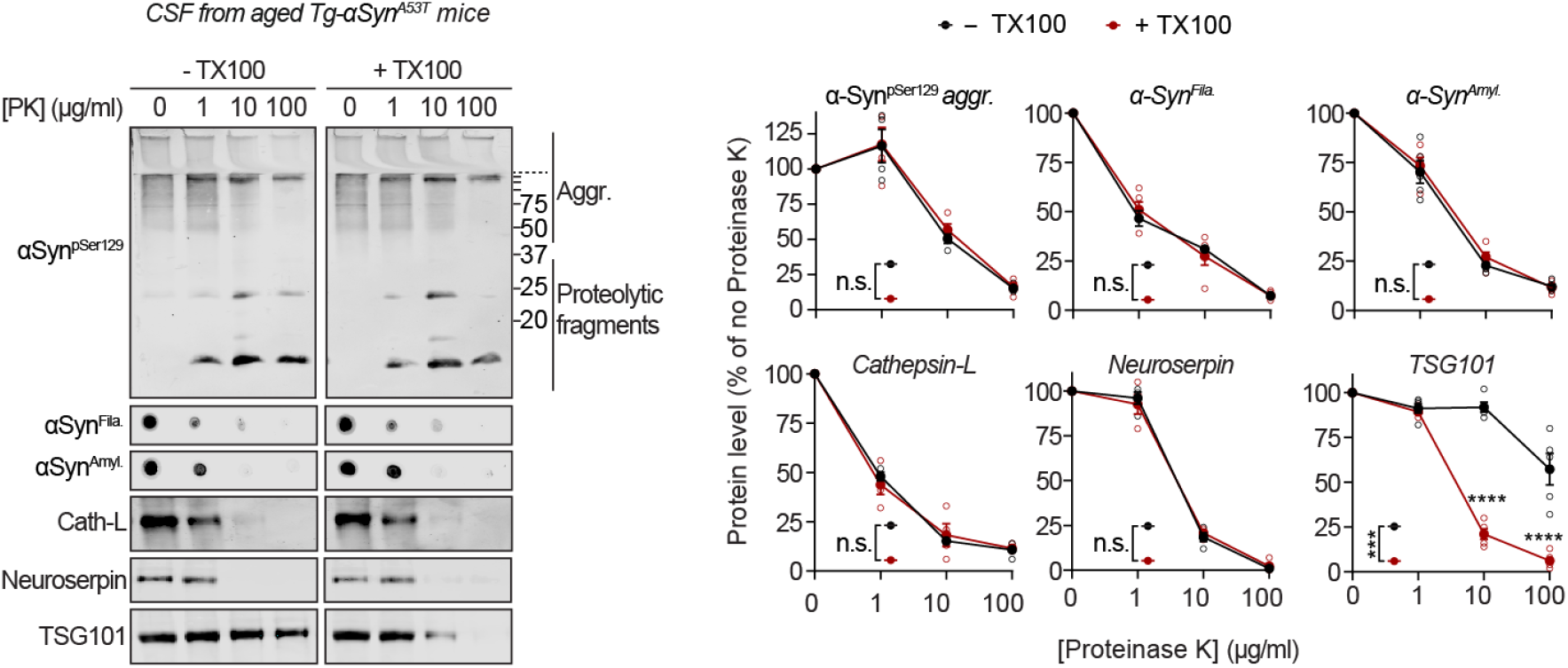
Extracellular αSyn species in cerebrospinal fluid are not membrane-enveloped. Cerebrospinal fluid (CSF) collected from 6 month old Tg-αSyn^A53T^ mice were subjected to limited proteolysis using indicated concentrations of proteinase K (PK) in the absence or presence of 0.1% Triton X-100. Immunoblots for αSyn protein levels (αSyn^pSer129^, αSyn^Amyl^, and αSyn^Fila^), cathepsin-L, TSG101, and neuroserpin (n=4). All data represent means ± SEM. Each ‘n’ is an independent proteolysis experiment. ***P<0.001; ****P<0.0001 by RM 2-way ANOVA and Bonferroni post-test with multiple hypothesis correction.

**Supplementary Figure S7.**
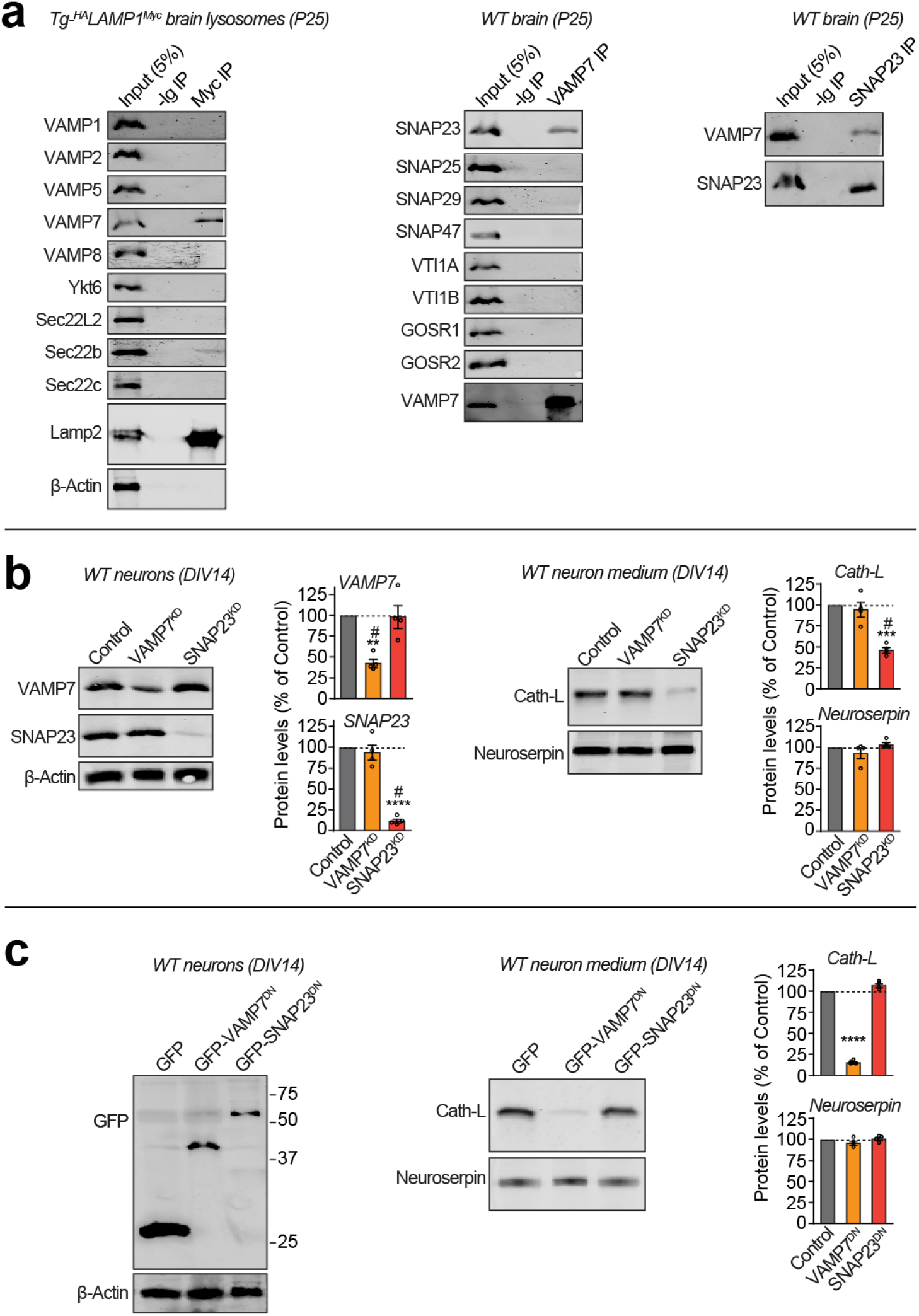
SNARE-dependence of lysosomal exocytosis. (**a**) From Tg-^HA^Lamp1^Myc^ mouse brains, we immunoisolated lysosomes (as in Fig. 2), followed by immunoblotting for the indicated vSNAREs (left panel). From wild type mouse brains, we immunoprecipitated (IP) VAMP7 and immunoblotted for the indicated co-IP’d Qb motif containing tSNAREs (middle panel). We then performed the inverse IP, where SNAP23 was IP’d and VAMP7 co-IP was tested by immunoblotting (right panel) (representative of n=3). (**b**) In WT neurons, shRNA knockdown constructs VAMP7^KD^ and SNAP23^KD^ were lentivirally expressed (Control = virus without shRNA), followed by quantitative immunoblotting of VAMP7 and SNAP23 levels (n=4). Secreted levels of lysosome luminal protein cathepsin-L and the constitutively secreted protein neuroserpin were also quantified from the medium, and normalized to β-actin quantified from the respective culture lysate (n=4). (**c**) In WT neurons, dominant-negative constructs GFP-VAMP7^KD^ and GFP-SNAP23^KD^ were lentivirally expressed (Control = GFP), followed by immunoblotting for GFP (representative of n=4). Quantification of cathepsin-L and neuroserpin in the medium, normalized to β-actin in cell lysates (n=4). All data represent means ± SEM. Each ‘n’ is an independent immunoisolation/immunoprecipitation in (**a**) and an independently infected neuron culture in (**b-c**). In (**b-c**) **P<0.01; ***P<0.001; ****P<0.0001 by RM 1-way ANOVA with Dunnett multiple-comparison correction; and ^#^P<0.05 by non-parametric Friedman test with Dunn’s multiple-comparison adjustment.

**Supplementary Figure S8.**
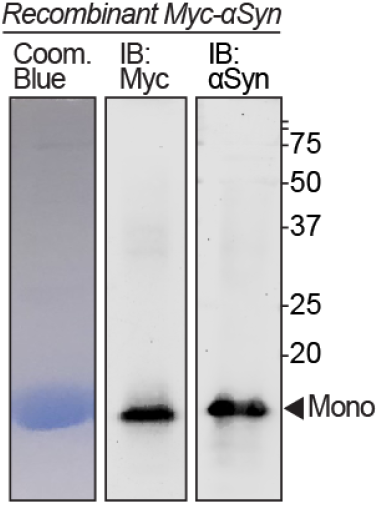
Recombinant myc-αSyn. Purified recombinant myc-αSyn protein separated by SDS-PAGE, followed by Coomassie brilliant blue staining, as well as immunoblotting against myc and αSyn.

## METHODS

### Mouse lines and husbandry

Mice were housed with a 12 h light/dark cycle in a temperature-controlled room with free access to water and food. Animal husbandry and the experimental protocols used in this study were approved by the Institutional Animal Care and Use Committee (IACUC) at Weill Cornell Medicine.

### CSPα knockout mice rescued by transgenic αSyn^A53T^ expression

Transgenic mice that express αSyn^A53T^ under the control of Thy-1 promoter (Chandra et al., 2005) were crossed to CSPα knockout mouse line (Fernández-Chacón et al., 2004) by breeding Tg-αSyn^A53T^ to CSPα^+/−^ mice. The early neurodegeneration and death of CSPα^−/−^ mice was rescued by overexpressing αSyn^A53T^ (Chandra et al., 2005). Rescued Tg-αSyn^A53T^/CSPα^−/−^ mice were bred with CSPα^+/−^ mice to obtain littermate CSPα^+/−^ and CSPα^−/−^ mice that either express or lack αSyn^A53T^ transgene. CSPα^+/−^ mice have no CSPα loss of function phenotype. Both, Tg-αSyn^A53T^ and CSPα^−/−^ mouse lines are available from Jackson Laboratory: B6.Cg-Tg(THY1-SNCA*A53T)M53Sud/J (stock # 008135) and B6.129S6-Dnajc5tm1Sud/J (stock # 006392), respectively.

### Generation of Tg-^HA^Lamp1^Myc^ mice

The transgenic mice were derived from a lentiviral vector. The lentiviral vector included truncated synapsin-1 promoter followed by ORF including Lamp1 signal sequence - 2xHA epitope tag – rat Lamp1 – 6His – TEV cleavage site - 2xMyc epitope tag. This lentivirus vector was digested overnight at 37°C with PacI and SphI to create a 3.45 kb linear fragment for pronuclear microinjection. The cleavage product included truncated 5’ LTR, the promoter and ^HA^Lamp1^Myc^ ORF (as described above), WPRE, and part of the bGH poly(A) signal. The enzymes were heat inactivated at 65°C for 20 minutes and the 3.45 kb band was purified/extracted using the Qiaquick gel extraction kit. The pronuclear injection was performed at Cornell University Stem Cell & Transgenic Core Facility into B6(Cg)-Tyrc-2J/JxFVB embryos. 115 embryos were injected, and 76 embryos proceeded to the 2-cell stage. Of the 18 clones born, 2 positive clones (male founders 1 and 7) contained PCR-detectable full length ^HA^Lamp1^Myc^, followed by confirmation by immunoblotting for protein expression. The founder transgenic clones were bred to wild type female C57BL/6 mice. The genotyping primers amplify sequences either near the N-terminal end (F: 5’-CGCGACCATCTGCGCTG-3’ and R: 5’-GCTGTGCCGTTGTTGTC-3’; product = 297 bp), or near the C-terminal end (F: 5’-GCACATCTTTGTCAGCAAGGCG-3’ and R: 5’-GCAATAGCATGATACAAAGGC-3: product = 474 bp) of the insert.

### *Crossing* Tg-αSyn^A53T^ *mice with Tg-^HA^Lamp1^Myc^ mice*

Neuron-specific ^HA^Lamp1^Myc^ transgenic mice were crossed to Tg-αSyn^A53T^ mice, and bred either as single transgenics (Tg-^HA^Lamp1^Myc^ to Tg-αSyn^A53T^) or bred as Tg-αSyn^A53T^/Tg-^HA^Lamp1^Myc^ double-transgenic to wild type. Either breeding-scheme produced Mendelian 25% Tg-αSyn^A53T^/Tg-^HA^Lamp1^Myc^ and 25% wild type progeny, and 25% progeny carrying each transgene alone; indicating single copy or insertion site for each transgene, carried on distinct chromosomes.

### Lysosome (dextranosome) isolation from mouse brains via density gradient centrifugation

Enrichment of dextran-loaded lysosomes (dextranosomes), enhanced by mitochondrial swelling, was based on previously published techniques (Arai et al., 1991; Graham, 2001). Mice were anesthetized with isoflurane and intracranially injected with 10 µl dextran-70 (250 mg/ml) into each cortex. 36 h later, mice were terminally anesthetized and perfused with homogenization buffer (HM = 10 mM HEPES buffer pH 7.0 with 0.25 M sucrose and 1 mM EDTA). The brain was dissected out, washed with HM and homogenized in 1.5 ml HM. Homogenate was centrifuged at 340 g_av_ for 5 min, any floating fat layer was aspirated, pellet discarded, and the supernatant was re-centrifuged at 340 g_av_ for 5 min. The resultant post-nuclear supernatant was incubated with 1 mM CaCl_2_ for 5 min at 37°C to swell the mitochondria, in order to reduce mitochondrial contamination in the dextranosomal fractions. This solution was centrifuged at 10,000 g_av_ for 30 min to precipitate heavy organelles. The pellet containing swollen mitochondria, peroxisomes and dextranosomes was resuspended in 1 ml HM and layered over 9 ml of 27% v/v Percoll in 0.25 M sucrose, followed by centrifugation at 35,000 g_av_ for 90 min, to generate the Percoll gradient. 1 ml fractions were collected from top, with dextranosomes expected near the bottom of the gradient. Fractions were centrifuged at 100,000 g_av_ at for 1 h to pellet the Percoll particles. The supernatant above the Percoll pellet was concentrated using Amicon (10 kDa cutoff), and assayed.

### Assays for cathepsin-D, citrate synthase, and catalase activity

Activities of enzymes contained within the lysosomes (cathepsin-D), mitochondria (citrate synthase), or peroxisomes (catalase) were measured from the Percoll gradient fractions dissolved in 0.1% Triton X-100 (final). Cathepsin-D activity was assayed using ab65302 kit (Abcam) according to the manufacturer’s protocol. Each fraction was incubated with the cathepsin-D substrate GKPILFFRLK(Dnp)-D-R-NH_2_ labeled with MCA at 37°C for 1 h in the dark. Fluorescence was measured at Ex: 328 nm, Em: 460 nm, in a solid white 96-well plate (Costar). Citrate synthase activity was assayed using ab239712 kit (Abcam) according to the manufacturer’s protocol, and absorbance at 412 nm was measured in clear bottom 96-well plate (Costar). Catalase activity was assayed using ab83464 kit (Abcam) according to the manufacturer’s protocol, and absorbance at 570 nm was measured in clear bottom 96-well plate (Costar). Synergy H1 Hybrid Reader was used for all three enzyme assays.

### Neuromuscular behavior tests and survival study

#### Pole test

The pole test was performed based on the method established by Ogawa et al. (Ogawa et al., 1985). Animals were positioned with their head upward near the top of a wooden dowel (1 cm in diameter and 50 cm high). The time taken until they turned completely downward (defined as a “T-turn”, indicative of bradykinesia) and time taken after the T-turn to descend to the bottom (indicative of motor coordination) were recorded. All animals were trained three times before performing the test. The maximum time allowed for each measurement was 20 s. If the animal could not turn, or fell during descent, the experiment was repeated; and if the animal again could not turn or fell during descent, the time for the activity was recorded as the maximum 20 s. *Grid hang test.* As in (Burré et al., 2010), animals were placed on top of a wire mesh grid (1.27 cm × 1.27 cm). The grid was then shaken lightly to cause the mouse to grip the wires with all four limbs, and then turned upside down. The mesh was held approximately 20 cm above the home cage litter, high enough to prevent the mouse of easily climbing down but not to cause harm in the event of a fall. A stopwatch was used to record the time taken by the animal to fall off the grid. Three trials per mouse were performed with a 1 min inter-trial interval. The highest time of the 3 trials was assigned as the time for each animal to fall. After a maximum hang time of 120 s, mice were removed from the grid. *Hind limb clasping.* Mice were lifted by tail for 20 s. If an animal retracted its hind limbs toward the abdomen and held them there, it was scored positive for hind limb clasping. *Survival study.* End point for survival study was reaching neuromuscular debility (e.g. paresis) which could make it difficult for the animal to get to food or water. At that time, the animal was euthanized according to the approved protocols.

### Immuno-isolation of lysosomes from mouse brains

Immunoprecipitation was used to isolate lysosomes from Tg-^HA^Lamp1^Myc^ mouse brains. Mouse brains were lysed in PBS without detergent and supplemented with EDTA-free protease inhibitor cocktail (Thermo Fisher). The post-nuclear supernatant was incubated with anti-myc magnetic beads (with covalently cross-linked clone 9E10 mAb; Pierce) for 2 h at 4°C, followed by 3 washes in PBS. The precipitated material was eluted in non-reducing 2x Laemmli sample buffer at room temperature, followed by immunoblotting.

### Mouse CSF collection and limited proteolysis

6 month old Tg-αSyn^A53T^ mice were terminally anesthetized with isoflurane, and CSF was collected by exposing cisterna magna (Lim et al., 2018), and using syringe aspiration (Hamilton 10 μl syringe). Samples with visible signs of blood were discarded. Following CSF collection, the mice were secondarily euthanized by cervical dislocation and brain collected for unrelated experiments. Due to low and/or variable volume of CSF collected in different attempts, multiple littermate mice were tapped and the CSF was pooled prior to proteolysis experiments.

Fresh pooled CSF at room temperature was aliquoted into 2 equal portions, and equal volume of either PBS or PBS containing Triton X-100 (0.1% final) was added to each tube and incubated for 10 min. Samples were then mixed with equal volume of Proteinase K (0, 1, 10, or 100 μg/ml final) from 100x stocks in sterile water, and incubated for 1 h at 15°C (to minimize membrane permeability). Proteolysis was immediately stopped by adding 5x Laemmli sample buffer with PMSF (1 mM final), followed by immunoblotting.

### Immunoprecipitation

For co-immunoprecipitation of SNARE proteins, brain homogenate was lysed in PBS containing 0.1% Triton X-100 and supplemented with EDTA-free protease inhibitor cocktail (ThermoFisher). Postnuclear supernatant was incubated for 2 h at 4 °C with the antibody, and then for another 1 h following the addition of protein-G Sepharose (GE Healthcare). Protein bound to antibody beads was washed five times with lysis buffer at 4 °C, and eluted in 2x Laemmli sample buffer. Eluent was boiled for 20 min to disassemble SNARE complexes, followed by immunoblotting.

### Long term primary neuron culture from Tg^2x^-αSyn^A53T^ mice

Primary cultures were based on modification of our neuron culture protocol (Naseri et al., 2020) with published procedures for long term cultures (Lesuisse and Martin, 2002). Neonatal (P0) cerebral cortices were isolated and dissected in cold HBSS. Cortices were dissociated by trypsinization (0.05% trypsin-EDTA for 10 min at 37 °C), triturated with a siliconized pipette and plated onto poly-l-lysine-coated 24-well plastic dishes at high density (800-1000 cells/mm^2^; each experiment had the same cell density across plates), in DMEM (Invitrogen) supplemented with 5% horse serum (Invitrogen) and 1% penicillin-streptomycin (Life Technologies). Humidity was enhanced by adding autoclaved water in spaces between the wells, and the plates were not disturbed for at least 5 days. On day 5-7, medium was changed to serum-free neurobasal medium (Life Technologies) supplemented with 2% B27 (Life Technologies) or N21-MAX (R&D Systems), and neurons were infected at the same time with lentiviruses. 50% of the medium was changed every 7 days. Neuronal cultures were maintained for up to 7 weeks (DIV 49).

Tg^2x^-αSyn^A53T^ mouse breeders were derived from Tg-αSyn^A53T^ mice crossed to each other. However, this line was difficult to breed due to short survival and unreliable breeding output. Alternatively, we cultured cortices from all of the pups from either Tg-αSyn^A53T^ parents, or from Tg^2x^-αSyn^A53T^ male bred with Tg-αSyn^A53T^ females (harem breeding). The pups were tested for homozygosity by cerebellar immunoblots against αSyn. Only Tg^2x^-αSyn^A53T^ cultures were maintained through to the DIV 5 steps of medium replacement and lentiviral infection.

Media from 49 day old neurons was spun through 5 um mini strainer (pluriselect) to exclude any debris, followed by concentration on 10 kDa Amicon (Millipore).

### Lentivirus production and transduction

HEK293T cells were co-transfected with the lentiviral vector plasmid, with the HIV-1 lentiviral packaging constructs pRSVREV and pMDLg/pRRE (Dull et al., 1998), and the vesicular stomatitis virus-G expression plasmid pHCMVG (Yee et al., 1994). The virus-containing culture supernatant was collected 48 h post-transfection and was concentrated by centrifugation at 50,000 gav for 90 min. The viral pellet was resuspended in neuronal medium (at 1/10 of the pre-centrifugation volume). All lentiviruses used in a single experiment were prepared together. Neurons were infected on DIV 5-7, and harvested for experiments at times described in figure captions.

### Immunofluorescence in neurons

For immunofluorescent labeling of cortical neuron cultures, cells on coverslips were washed with PBS + 1 mM MgCl2 and fixed in 4% paraformaldehyde for 30 min at room temperature. Cells were permeabilized for 5 min in PBS + 0.1% Triton X-100, and blocked for 20 min in PBS + 5% BSA (blocking buffer). Coverslips were then incubated in primary antibodies in blocking buffer overnight at 4 °C. Following five washes, coverslips were incubated with secondary antibodies labeled with Alexa 488 and/or Alexa 546 (Life Technologies) in blocking buffer containing DAPI, for 1–2 h, followed by five PBS washes and mounting on slides in Fluormount G (ThermoFisher). Images were acquired on a Nikon H550L microscope.

### SNARE shRNA knockdown and dominant-negative strategies

Once VAMP7 and SNAP23 were identified as cognate lysosomal SNAREs, in agreement with prior studies (Rao et al., 2004), we modulated their function via shRNA knockdown and dominant-negative strategies: Based on pre-characterized target sequences and hairpins described on the Broad Institute Genetic Perturbation Platform, mouse VAMP7 and SNAP23 shRNAs were cloned into L309 lentiviral vector to be expressed under human H1 promoter. For VAMP7, the target sequence was 5’-TTACGGTTCAAGAGCACAAAC-3’, and for SNAP23, the target sequence was 5’-CAACCGAGCCG-GATTACAAAT-3’. Rescue expression of human SNAP23 from FUW vector (Lois et al., 2002) was not affected by the anti-mouse shRNA target sequence.

To express a dominant negative VAMP7 fragment, rat VAMP7/TI-VAMP truncated at the C-terminal end (Ala^2^-Asn^120^ fragment expressed) (Martinez-Arca et al., 2000) was expressed as an eGFP chimera (GFP-VAMP7^2-120^) via FUGW lentiviral vector (Lois et al., 2002).

To generate a dominant negative version of SNAP23, the last eight residues of human SNAP23 were deleted (Asp^2^-Ala^203^ segment expressed) mimicking the dominant negative SNAP25^1-197^ fragment generated by Botulinum toxin-A cleavage (Huang et al., 1998) and expressed as an eGFP chimera (GFP-SNAP23^2-203^) from FUGW lentiviral vector (Lois et al., 2002).

### Proximity-labeling of lysosomal contents

To target Apex-2 to the lysosomal lumen, Apex-2 was fused at the N-terminus to the transmembrane domain of rat Lamp1. The ^Apex-2^Lamp1 construct consists of: Lamp1 signal sequence - Flag tag - APEX2 - rat Lamp1 truncation including transmembrane domain plus C-terminal cytosolic tail with lysosome-targeting motif (Lamp1^370-407^ including the targeting motif ^403^GYQTI^407^) - 6His - TEV - Myc tag, generating a 36 kDa protein.

^Apex-2^Lamp1 was lentivirally expressed via a truncated synapsin-1 promoter in Tg^x2^-αSyn^A53T^ primary neurons by infection on DIV7. On DIV 49, cells were incubated at 37°C for 1 h in medium containing 500 μM biotin tyramide, plus a lysosomal pH-raising cocktail (10 nM NH^4^Cl and 100 nM Bafilomycin A1) to enhance the reaction. H_2_O_2_ was added at a final concentration of 1 mM for exactly 1 min at room temperature. Reaction was quenched for 30 sec with PBS + 1mM MgCl_2_ containing antioxidants (5 mM Trolox, 10 mM ascorbic acid). Cells were washed with PBS + 1mM MgCl_2_, and lysed in PBS + 0.1% Triton-X 100, and biotinylated proteins were precipitated on streptavidin-magnetic beads (Dynabeads; Thermo), with 3 washes before elution in 2x Laemmli sample buffer.

To chase biotin-labeled proteins, Tg^x2^-αSyn^A53T^ primary neurons lentivirally expressing ^Apex-2^Lamp1, were subjected to biotin labeling on DIV 47, similar to above, except using 0.5 mM H_2_O_2_ to reduce toxicity. The reaction was quenched with 37°C medium containing antioxidants (5 mM Trolox, 10 mM sodium ascorbate, and 5 mM glutathione). 10 minutes later, the medium was replaced, containing 2.5 mM glutathione. 48 h later, the culture-medium was collected, and biotinylated proteins were precipitated on streptavidin-magnetic beads (Dynabeads; Thermo), with 3 washes before elution in 2x Laemmli sample buffer.

### Purification and in vitro aggregation of recombinant myc-αSyn

Recombinant myc-αSyn was expressed and purified essentially as in (Burré et al., 2015). Full-length human αSyn cDNA containing an N-terminal myc epitope-tag was inserted into a modified pGEX-KG vector (GE Healthcare), after a TEV protease recognition site to allow cleavage from GST (myc-αSyn contains an extra N-terminal glycine after cleavage with TEV protease). BL21 strain of *E. coli* bacteria transformed with this plasmid were grown to optical density 0.6 (at 600 nm), and protein expression was induced with 0.05 mM isopropyl β-D-thiogalactoside (IPTG) for 6 hours at room temperature. Bacteria were pelleted by centrifugation for 20 min at 4000 rpm, and then resuspended in solubilization/lysis buffer, PBS containing 0.5 mg/ml lysozyme, 1 mM PMSF, DNase, and an EDTA-free protease inhibitor mixture (Roche). Bacteria were further broken by sonication, and insoluble material was removed by centrifugation for 30 min at 7000 g_av_ at 4°C. Protein was affinity-bound to glutathione Sepharose beads (GE Healthcare) by incubation overnight at 4°C, followed by TEV protease (Invitrogen) cleavage for 6 hours at room temperature. The His-tagged TEV protease was removed by incubation with Ni-NTA (Qiagen) overnight at 4°C. Protein concentration was assessed using the bicinchoninic acid (BCA) assay (Thermo Fisher Scientific).

For the aggregation studies, 10 μl of concentrated extracellular medium from neuronal cultures – collected over 7 days (DIV 42-49), spun through 5 μm strainer (Puriselect), and concentrated 10x using 10 kDa cutoff Amicon (Millipore) – was added to 40 μl of recombinant myc-αSyn (to 4 μg/μl final concentration) in PBS with protease inhibitors. This mixture was incubated at 37°C with shaking at 300 rpm, while sample aliquot were taken at indicated time periods and frozen at −80°C. At the end of final incubation time-point, all the samples were thawed and measured together.

5 μl of sample was mixed with 95 μl of 100 μM K114 (Santa Cruz Biotechnology) in 100 mM glycine-NaOH, pH 8.45, and K114 fluorescence was measured at Ex: 390 nm, Em: 535 nm, in a solid white 96-well plate (Costar) using a plate reader (Synergy H1 Hybrid Reader, BioTek). Thioflavin-T measurement was done similarly, by mixing 5 μl sample with 95 μl of 25 μM Thioflavin-T (Sigma) in PBS, pH 7.4, and fluorescence was measured at Ex: 450 nm, Em: 485 nm. After fluorometry, the same samples were dissolved in Laemmli sample buffer and used for SDS-PAGE or dot blots, followed by immunoblotting.

### SDS-PAGE and quantitative immunoblotting

For SDS-PAGE, 10-15% Laemmli gels (10.3%T and 3.3%C) were used to separate proteins on Bio-Rad apparati. 1 mM final DTT was used in SDS Laemmli buffer. Proteins were transferred onto nitrocellulose (pore-size = 0.45 μm; GE Healthcare) and blocked with 5% w/v fat-free dry milk in tris buffered saline, pH 7.5 supplemented with 0.1% Tween 20 (TBS-T). Immunoblotting was performed by incubating the blocked membranes with primary antibodies in blocking buffer for 8-16 hours. Following 5 washes with TBS-Tween 20 (0.1%). Blots were incubated with secondary antibodies (goat anti-rabbit conjugated to IRDye 600RD or 800CW; LI-COR) at 1/5000 in blocking buffer for 1-3 hours. Immunoblots were washed 5x with TBS-T and dried, then scanned on Odyssey CLx (LI-COR) and quantified using Image-Studio software (LI-COR).

For ATP5G immunoblots, following the transfer, nitrocellulose membranes were dried and fixed for 15 min at room temperature in 1% paraformaldehyde in PBS (Ikegaki and Kennett, 1989). Membranes were then washed 3x with TBS-T and treated as above.

For dot blots, samples were dotted onto dry nitrocellulose membrane (if sample volume <5 μl) or on a PBS wetted nitrocellulose membrane under vacuum (if volume was >5 μl), and allowed to dry. The membrane was then immunoblotted as above.

### Antibody list

β-Actin: Sigma (A1978); A11 oligomers: Stressmarq (SPC-506D); ATP5G: Abcam (ab181243); Calreticulin: Thermo Fisher (OTI15F5) and Novus (NB600-103); Cathepsin-L: Novus (JM10-78); CD81: Novus (SN206-01); EEA1: Thermo Fisher (MA5-14794); Flag: Sigma Cl. M2 (F3165); GAPDH: Cell Signaling Cl. 14C10 (2118); GFP: Takara Cl. JL8 (632381) and Invitrogen (A11122); GluN2B/NR2B: Cell Signaling Cl. D15B3 (4212); GOSR1: Abcam (ab53288); GOSR2: ProteinTech (12095-1-AP); HA epitope: Abcam Cl. 16B12 HA.11 (ab130275); Histone H3: Cell Signaling Cl. 96C10 (3638); Hsp60: Abcam (ab5479); Lamp-1: DSHB Cl. H4A3 and Cl. 1D4B DSHB Rat mono; Lamp-2: Cl. ABL-93 DSHB Rat mono; Map-2: Millipore (AB5622); Myc epitope: DSHB Cl. 9E10 and Sigma (C3956); Na/K-ATPase: DSHB Cl. A6F; NeuN: Millipore Cl. A60 (Mab377); Neuroserpin: Abcam (ab33077); Pex3: Thermo Fisher (PA5-37012); Pex13: Sigma (ABC143); Sec22b: SYSY (186 003); Sec22c: Novus (NBP1-30760); Sec22L2: MyBiosource.com (MBS8507611); SNAP23: Santa Cruz (sc-166244) and Sudhof lab P914; SNAP25: Biolegend Cl. SMI81 (836304) and Sudhof lab P913; SNAP29: SYSY (111303); SNAP47: SYSY (111403); synaptogyrin-1: SYSY (103 002); αSyn: BD Transduction (610787) and Sudhof Lab T2270; αSyn filamentous (αSyn^Fila^): Abcam Cl. MJFR-14-6-4-2 (ab209538); αSynuclein Ser129 phosphorylated (αSyn^pSer129^): Abcam (ab51253); TGN38: BD Transduction (610899); TIM23: BD Transduction (611223); TSG101: Abcam (ab125011); α-Tubulin: DSHB Cl. 12G10; VAMP1: SYSY (104 002); VAMP2: SYSY (104 202); VAMP5: SYSY (176 003); VAMP7/TiVAMP: SYSY (232 011) and SYSY (232 003); VAMP8: SYSY (104 302); VTI1A: SYSY (165 002); VTI1B: SYSY (164 002); Ykt6: Abcam (ab241382).

### Statistical analyses

Each “n” consisted of reagents produced in parallel (e.g., purified proteins, lentivirus production, culturing neurons, brain or CSF harvested from mice euthanized together) and experiments performed in parallel (e.g. lentiviral infection, sample collection for immunoprecipitation and immunoblotting, mouse behavior recorded on same day etc.). Experiments were quantified using parametric as well as non-parametric statistical tests when warranted (see the tests described below and in figure captions). Randomization and coding of mice or samples was done by different investigator(s) than the investigator who did quantification and analysis. All analyses are performed using GraphPad Prism 8 and described in the figure legends. Column analyses of data with more than 2 groups depicted in bar graphs (e.g. αSyn species in lysosomal fractions and αSyn species in extracellular media) were analyzed by repeated measures (RM) 1-way ANOVA with Dunnett multiple-comparison correction, as well as using the non-parametric Friedman test with Dunn’s multiple-comparison adjustment. For comparison of only two groups (e.g. such as for littermate ratio of WT and Tg-^HA^Lamp1^Myc^ cross) 2-tailed Student’s t-test was used. For time-course experiments (e.g. accumulation of αSyn over age and seeding of recombinant αSyn) the curves were analyzed using RM 2-way ANOVA in comparing the overall curve, plus the Bonferroni post-test with multiple hypothesis correction to compare data at each time-point independently or to examine more than two groups. Graphs depicting the correlation (e.g. between the levels of pathogenic αSyn species released versus the levels of cathepsin-L released in the medium) were analyzed via Pearson’s correlation and shown with linear regression lines, as well as the 95% confidence interval using dotted lines. Kaplan-Meier curves (e.g. survival, onset of hind-limb clasping, and onset of grid-hang impairments) were compared via Log-rank (Mantel-Cox) test. For vertical pole studies, due to decreasing sample size with each mouse death, two separate curves were graphed to represent each of the pole-turn and the pole-descent data. Mice that died were scored as 20 s (maximum measurement) for rest of the trial – not as an actual measurement, but as a place holder. Data with 20 s placeholder were analyzed by RM 2-way ANOVA and data without the 20 s placeholder were analyzed via mixed effects analysis.

## ACKNOWLEDGEMENTS

We thank Dr. Tom Südhof for kindly sharing the CSPα knockout and Tg-αSyn^A53T^ mouse lines, as well as antibodies against αSyn, SNAP-23, and SNAP-25.

## FUNDING

This work was supported by NIH F31 studentship (NS098623, to N.N.N.), grants from Alzheimer’s Association (NIRG-15-363678 to M.S.), American Federation for Aging Research (New Investigator in Alzheimer’s Research Grant, to M.S.), NIH National Institute for Neurological Disorders and Stroke (1R01NS102181 and 1R01NS113960, to J.B.; and 1R01NS095988, to M.S.), NIH National Institute for Aging (1R01AG052505, to M.S.).

## AUTHOR CONTRIBUTIONS

Y.X.X., N.N.N., J.F., P.K., Y.N., J.B., and M.S. designed, performed and analyzed all experiments. Y.X.X., N.N.N., J.B., and M.S. wrote the manuscript. M.S. conceived the project and directed the research.

## COMPETING INTERESTS

The authors declare no competing interests.

## Notes

### Competing Interest Statement

The authors have declared no competing interest.

